# Oxygen chemoreceptor inhibition by dopamine D_2_ receptors in isolated zebrafish gills

**DOI:** 10.1101/2024.10.08.617247

**Authors:** Maddison Reed, Michael G. Jonz

## Abstract

Dopamine is an essential modulator of oxygen sensing and control of ventilation and was the first neurotransmitter described in the carotid body. Little is known of the evolutionary significance of dopamine in oxygen sensing, or whether it plays a similar role in anamniotes. In the model vertebrate, zebrafish (*Danio rerio*), presynaptic dopamine D_2_ receptor expression was demonstrated in gill neuroepithelial cells (NECs)—analogues of mammalian oxygen chemoreceptors; however, a mechanism for dopamine and D_2_ in oxygen sensing in the gills had not been defined. The present study tested the hypothesis that presynaptic D_2_ receptors provide a feedback mechanism that attenuates the chemoreceptor response to hypoxia. Using an isolated gill preparation from Tg(*elavl3*:GCaMP6s) zebrafish, we measured hypoxia-induced changes in intracellular Ca^2+^ concentration ([Ca^2+^]_i_) in NECs and postsynaptic neurons. Activation of D_2_ with dopamine or specific D_2_ agonist, quinpirole, decreased hypoxic responses in NECs; whereas D_2_ antagonist, domperidone, had the opposite effect. Addition of SQ22536, an adenylyl cyclase (AC) inhibitor, decreased the effect of hypoxia on [Ca^2+^]_i_, similar to dopamine. Activation of AC by forskolin partially recovered the suppressive effect of dopamine on the Ca^2+^ response to hypoxia. Further, we demonstrate that the response to hypoxia in postsynaptic sensory neurons was dependent upon innervation with NECs, and was subject to modulation by activation of presynaptic D_2_. Our results provide the first evidence of neurotransmission of the hypoxic signal at the NEC-nerve synapse in the gill and suggest that a presynaptic, modulatory role for dopamine in oxygen sensing arose early in vertebrate evolution.

**Key points:**

1. For the first time, we present an experimental model that permits imaging of intracellular Ca^2+^ in identified vertebrate oxygen chemoreceptors using GCaMP in a whole/intact sensing organ.
2. The hypoxic response of zebrafish chemoreceptors is attenuated by dopamine through a mechanism involving D_2_ receptors and adenylyl cyclase.
3. Zebrafish oxygen chemoreceptors send a hypoxic signal to postsynaptic (sensory) neurons.
4. Postsynaptic neuronal responses to hypoxia are modulated by presynaptic D_2_ receptors, suggesting a link between chemoreceptor inhibition by dopamine and modulation of the hypoxic ventilatory response.
5. Our results suggests that a modulatory role for dopamine in oxygen sensing arose early in vertebrate evolution.

## Introduction

Dopamine is an important neuromodulator involved in oxygen sensing and control of reflex hyperventilation. In the mammalian carotid body, it is the most abundant neurotransmitter and has been described in numerous species (Gauda, 2002). Hypoxic activation of carotid body oxygen chemosensory type 1 (glomus) cells involves Ca^2+^-dependent neurotransmitter release to act on sensory terminals of the carotid sinus nerve (González et al., 1994; Kumar & Prabhakar, 2012; López-Barneo et al., 1988; Nurse, 2010). Dopamine, released by type 1 cells, has been shown to have autocrine–paracrine actions in the carotid body via G-protein-coupled dopamine D_2_ receptors on both presynaptic type 1 cells and postsynaptic afferent terminals (Nurse, 2010; Zhang et al., 2018, Carroll et al., 2005; Gauda, 2002; Itturiaga et al., 2009; Mir et al., 1984). At the presynaptic type 1 cell, the inhibitory actions of dopamine decrease further neurotransmitter release, thereby mediating carotid body hypoxia signaling (Benot & López-Barneo, 1990). Dopamine has also been implicated in regulation of chemosensitivity during acclimatization to prolonged periods of hypoxia (Bisgard, 2000; Huey & Powell 2000; Powell, 2007), and as an important factor in development or maturation of the carotid body (Carroll et al., 2005; De Caro et al., 2013; Gauda, 1996, 2002).

In fish, oxygen sensing occurs in the gills via chemoreceptive neuroepithelial cells (NECs) in a manner similar to carotid body type I cells. NECs are found in the gill filament epithelium and respond to a decrease in PO_2_ by inhibition of background K^+^ channels, membrane depolarization, and Ca^2+^-dependent vesicular recycling (Jonz et al., 2004; Zachar et al., 2017). The latter is consistent with neurotransmitter release into the synaptic cleft, which may lead to activation of sensory nerve fibers to control ventilation. NECs are polymodal, such that they are sensitive to changes in O_2_, CO_2_, H^+^ and lactate (Abdallah et al., 2015; Jonz, 2018; Qin et al., 2010; Leonard et al., 2022) and are characterized by immunoreactivity to serotonin (5-hydroxytryptamine [5-HT]) and synaptic vesicle protein, SV2 (Jonz & Nurse, 2003). Teleost gill arches are innervated by branches of the glossopharyngeal (IX) and vagus (X) cranial nerves, which carry parasympathetic efferent fibers to the gill vasculature and sensory (afferent) fibers from chemoreceptors of the gill filaments (Nilsson, 1984; de Graaf, 1990; Sundin and Nilsson, 2002). Beneath the efferent filament artery lies a series of 4-6 neuronal somata found along a nerve bundle that travels the length of the filament. These are referred to as chain neurons (ChNs) and may provide an additional source of sensory innervation to NECs (Jonz & Nurse, 2003).

The gill arches in fish are homologues of the sites of O_2_-sensing organs in mammals, such as the carotid body and pulmonary neuroepithelial bodies; however, whether gill NECs are functionally similar (i.e. homologous) with carotid body type I cells or neuroepithelial bodies, is controversial (Milsom & Burleson, 2007; Zachar & Jonz, 2012, Hockman et al., 2017; Jonz, 2024). Despite the similarities between NECs and type I cells, and in contrast to the well-described neurochemistry of the carotid body (Nurse, 2010), a mechanism for control of ventilation during hypoxia via neurotransmission in the gills of fish has never been defined.

Early evidence of a role for dopamine in the gills was shown in isolated gills of rainbow trout (*Oncorhynchus mykiss*), where dopamine caused a small, brief burst in afferent nerve activity followed by mild inhibition (Burleson & Milsom, 1995). In live, whole-animal experiments using zebrafish larvae, application of dopamine or the dopamine D_2_-receptor agonist, quinpirole, both decreased ventilation frequency, suggesting inhibitory effects of dopamine on ventilation (Shakarchi et al., 2013, Reed et al., 2024). In isolated gill tissue from adult zebrafish, quantitative polymerase chain reaction analysis confirmed expression of *drd2a* and *drd2b* (genes encoding D_2_ receptors) in the gills, and their relative abundance decreased following acclimation to hypoxia for 48 h (Reed et al., 2024). Further, immunohistochemical labelling in the gill localized D_2_ receptors to presynaptic NECs, as well as provided evidence for the synthesis and storage of dopamine by nerve terminals of postsynaptic sensory neurons that innervate NECs (Reed et al., 2024). While these findings point to an inhibitory role for dopamine via D_2_ receptors in the gill, a direct mechanism for how dopamine may participate in oxygen sensing has not been elucidated.

The goal of the present study was to delineate a mechanism by which presynaptic D_2_ receptors provide a feedback mechanism that attenuates the chemoreceptor response to hypoxia. Using a genetically-encoded Ca^2+^ indicator in gills isolated from transgenic zebrafish, we found that activation of presynaptic D_2_ receptors decreased the NEC Ca^2+^ response to hypoxia via intracellular adenylyl cyclase (AC) activation. Further, we provide evidence for postsynaptic modulation of the hypoxic signal in sensory neurons originating via activation of presynaptic D_2_ receptors.

## Methods

### Ethical approval

All wild-type and transgenic zebrafish were bred and maintained at the Laboratory for the Physiology and Genetics of Aquatic Organisms, University of Ottawa. Zebrafish were kept at 28°C on a 14:10-h light:dark cycle (Westerfield, 2007). Embryos and larvae were reared on a diet of rotifers and Gemma 75 (Skretting Canada, St. Andrews, NB), juveniles were fed Artemia and Gemma 150-300 (Skretting), and adults were fed Adult Zebrafish Diet (Ziegler Feeds, Gardners, PA, USA). Adult zebrafish were euthanized by concussion and decapitated. All procedures for animal use and euthanasia were carried out in accordance with institutional guidelines according to protocol BL-3666, and guidelines provided by the Canadian Council on Animal Care. The present work complies with the ethical guidelines set out by the journal and with those according to Animals in Research: Reporting In Vivo Experiments (ARRIVE).

In this study we used transgenic zebrafish Tg(*elavl3*:GCaMP6s) expressing the genetically encoded Ca^2+^ indicator GCaMP6s under the pan-neuronal promotor *elavl3* (Dunn et al. 2016). GCaMP6s contains the green fluorescent protein (GFP) as part of its structure. The Tg(*dat:tom20 MLS-mCherry*) line has previously been used to visualize dopaminergic neurons (Reed et al., 2024). In this line, the regulatory elements of the dopamine transporter gene (*dat*) were targeted to a reporter, mCherry, after fusion with the mitochondrial localizing signal (MLS) of Tom20 (Noble et al., 2015). To determine the distribution of dopaminergic nerve terminals relative to GFP-positive NECs, adult Tg(*elavl3*:GCaMP6s) fish were crossed with adult Tg(*dat:tom20 MLS-mCherry*) to generate double transgenic offspring containing both mCherry and GFP.

### Immunohistochemistry

Techniques for tissue extraction and immunolabeling were carried out as previously described (Jonz & Nurse, 2003). Whole gill baskets were removed and immersed in phosphate buffered solution (PBS) containing (mM): NaCl 137, Na_2_HPO_4_ 15.2, KCl 2.7, and KH_2_PO_4_ 1.5 at pH 7.8 (Bradford et al., 1994). Gill baskets were fixed by immersion in 4% paraformaldehyde in PBS overnight at 4°C. Tissues were removed and rinsed in PBS three times at 3 min before permeabilization for 24 h at 4°C. Permeabilizing solution (PBS-TX) contained 0.5-2% Triton X-100 in PBS (pH 7.8). After 3 rinses in PBS, gill baskets were then separated into individual arches. Gill arches were incubated in primary antibodies for 24 h at 4°C, rinsed with PBS three times at 3 min, and immersed in secondary antibodies for 1 h at room temperature in darkness.

NECs were identified using antibodies against serotonin (5-HT) or synaptic vesicle protein SV2. Polyclonal anti-5-HT was raised in rabbit against a 5-HT creatinine sulfate complex conjugated with bovine serum albumin (manufacturer specifications; cat. no. S5545, Sigma-Aldrich, Oakville, ON, Canada; Antibody Registry ID: AB_477522). Anti-5HT was used at 1:250 and localized with goat anti-rabbit secondary antibodies conjugated with fluorescein isothiocyanate (FITC, 1:50, cat. no. 111-095-003, Cedarlane, Burlington, ON, Canada). Monoclonal SV2 raised in mouse (AB_2315387 and AB_531908; Developmental Studies Hybridoma Bank, University of Iowa, IA, USA) was used at 1:100 and targeted by goat anti-mouse secondary antibodies conjugated with Alexa 594 at 1:100 (cat. no. A11005, Invitrogen, Burlington, ON, Canada). Neurons and nerve fibres were identified using antibodies against a zebrafish-specific neuronal marker (zn-12). Monoclonal anti-zn-12 raised in mouse (RRID: AB_2315387; Developmental Studies Hybridoma Bank, University of Iowa) was used at 1:100 and visualized using goat anti-mouse secondary antibodies conjugated with Alexa 594 at 1:100 (cat. no. A11005, Invitrogen, Burlington, ON, Canada). Labelling by these antibodies in the zebrafish gill has been previously characterized (Jonz & Nurse, 2003).

### Relative [Ca^2+^]_i_ measurements

Whole gill baskets were removed and separated into individual gill arches and immersed in extracellular solution containing (mM): 120 NaCl, 5 KCl, 2.5 CaCl_2_, 2 MgCl_2_, 10 HEPES, 10 glucose at pH 7.8. Isolated intact gill arches from Tg(*elavl3*:GCaMP6s) zebrafish were secured in a Petri dish using a metal tissue anchor (cat. no. 640251, Warner Instruments) and continuously perfused with extracellular solution (ECS) at pH 7.8. GFP-fluorescing cells were observed using a Nikon 40× water-immersion objective. NECs were identified by their size, presence of GFP, position along the centre of the filament epithelium, and location at the distal end of the filament. Using a Lambda DG-5 wavelength changer (Sutter Instruments, Novato, CA, USA), the preparation was exposed to 490 nm excitation light for 600 ms at a sampling frequency of 1 s^-1^. Images were captured with a CCD camera (QImaging, Surrey, BC, Canada), and fluorescence intensity was recorded with NIS Elements software (Nikon).

Dual recordings of NECs and chain neurons (ChNs) were carried out as above, with a modified depth of focus. Since these two cell types are located in different layers of the gill filament, the recording plane was set at an intermediate depth between the NEC and ChN, where both cell types were slightly out of focus, but still within the field of view, and therefore could be recorded simultaneously (see Fig. 9C).

### Solutions and drug treatments

High K^+^ extracellular solution was prepared with 90 mM NaCl, 35 mM KCl, 2.5 mM CaCl_2_, 2 mM MgCl_2_, 10 mM HEPES, 10 mM glucose. Zero Ca^2+^ extracellular solution was prepared with 120 mM NaCl, 5 mM KCl, 4.5 mM MgCl_2_, 10 mM HEPES, 10 mM glucose, and 1 mM EGTA. pH of all solutions was kept at 7.8. 100% N_2_ was bubbled through an air stone into solution reservoirs to create hypoxic solutions with a PO_2_ of approximately 25 mmHg. All control solutions were bubbled with compressed air for the same duration of time.

Drugs were introduced into the recording chamber by perfusion in extracellular solution. Nifedipine (cat. no. N7643, Sigma-Aldrich) and dantrolene (cat. no. 0507, Tocris) were used to block entry of extracellular Ca^2+^ and release of stored Ca^2+^, respectively. To target dopamine receptors, the D_2_ antagonist, domperidone (cat. no. D122; Sigma-Aldrich), and the D_2_ agonist, quinpirole (cat. no. Q102, Sigma-Aldrich), were tested. To identify an intracellular mechanism for D_2_-receptors, the AC inhibitor SQ22536 (cat. no 1435, Tocris) and AC activator forskolin (cat. no. 11018, Cayman Chemical) were tested. All drugs were first dissolved in dimethyl sulfoxide (DMSO) to produce a final DMSO concentration of <0.1%. At this concentration, DMSO had no effect on Ca^2+^ baseline and did not produce changes in fluorescence intensity (relative [Ca^2+^]_i_).

### Statistical Analysis

Throughout this study, hypoxic responses from a total of 73 cells were recorded from 41 adult zebrafish. For each recording, the baseline fluorescence was calculated as the average fluorescence intensity of a cell for the first 30 s in normoxia. All fluorescence values were divided by the baseline to evaluate changes in fluorescence intensity over time throughout a single recording. Statistical analysis was carried out using the Kruskal-Wallis test and Dunn’s test for multiple comparisons with Prism software (GraphPad Software Inc., La Jolla, CA, USA). All data were expressed as means ± standard error of the mean (SEM).

## Results

### NECs contain GCaMP6s and display a Ca^2+^ response to hypoxia *in situ*

The transgenic Tg(*elavl3*:GCaMP6s) zebrafish used in this study expresses a genetically-encoded Ca^2+^ reporter, GCaMP, under the control of the pan-neuronal promotor, *elavl3* (Dunn et al. 2016). To confirm expression of GCaMP in chemoreceptors, we used immunohistochemistry to identify GFP (part of the GCaMP complex) with known markers of gill NECs. Labelling of GFP-positive GCaMP-containing cells colocalized with anti-SV2 (Fig. 1A-C) and anti-5-HT (Fig. 1D-F), demonstrating that GCaMP is contained within NECs. We developed a procedure to record brief elevations in [Ca^2+^]_i_ that were associated with a hypoxic stimulus in single chemoreceptors *in situ* using isolated gills (Fig. 2). GCaMP-positive NECs were first identified using brightfield and 490 nm illumination (Fig. 2B), and then confirmed after successful Ca^2+^-imaging experiments by fixation and immunohistochemistry (Fig. 2C). These GCaMP-positive NECs displayed a Ca^2+^ response to hypoxia (Fig. 2D) and the magnitude of these responses did not change over time, or after multiple exposures (Fig. 2E, Kruskal-Wallis test, P>0.999, n=7).

**Figure 1.**
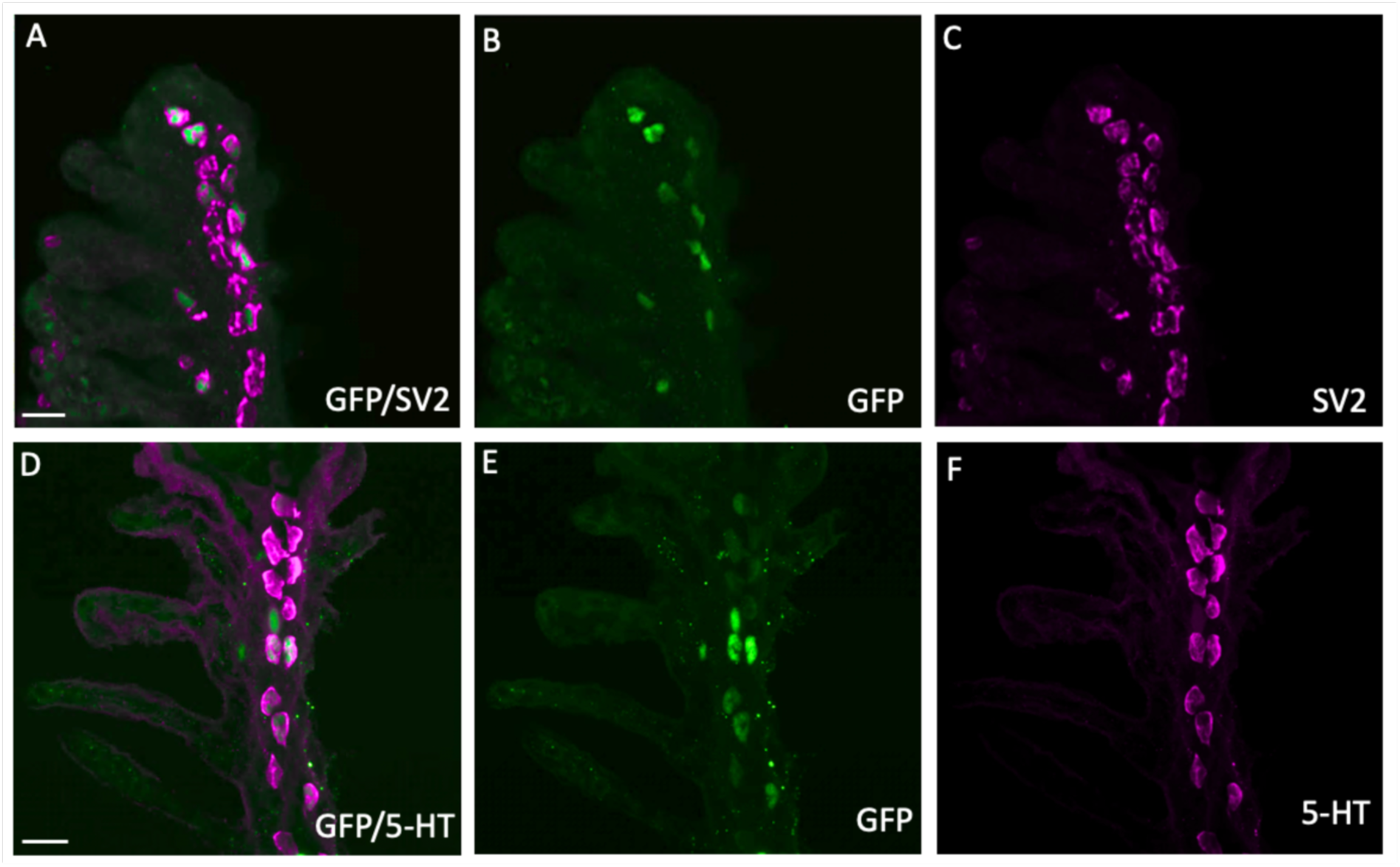
Characterization of GCaMP-positive neuroepithelial cells (NECs) in the gill epithelium of transgenic *elavl3*:GCaMP6s zebrafish. Confocal imaging of immunohistochemical localization of GCaMP with NECs containing synaptic vesicle protein-2 (SV2) and 5-hydroxytryptamine (5-HT). (A) Labelling with GFP (green) co-localized with NECs containing SV2 (magenta). (B, C) GCaMP and SV2 labelling shown separately. Scale bar in A=20 µm and applies to B and C. (D) Labelling with GCaMP (green) co-localized with NECs containing 5-HT (magenta). (E, F) GCaMP and 5-HT labelling shown separately. Scale bar in D=20 µm and applies to E and F.

**Figure 2.**
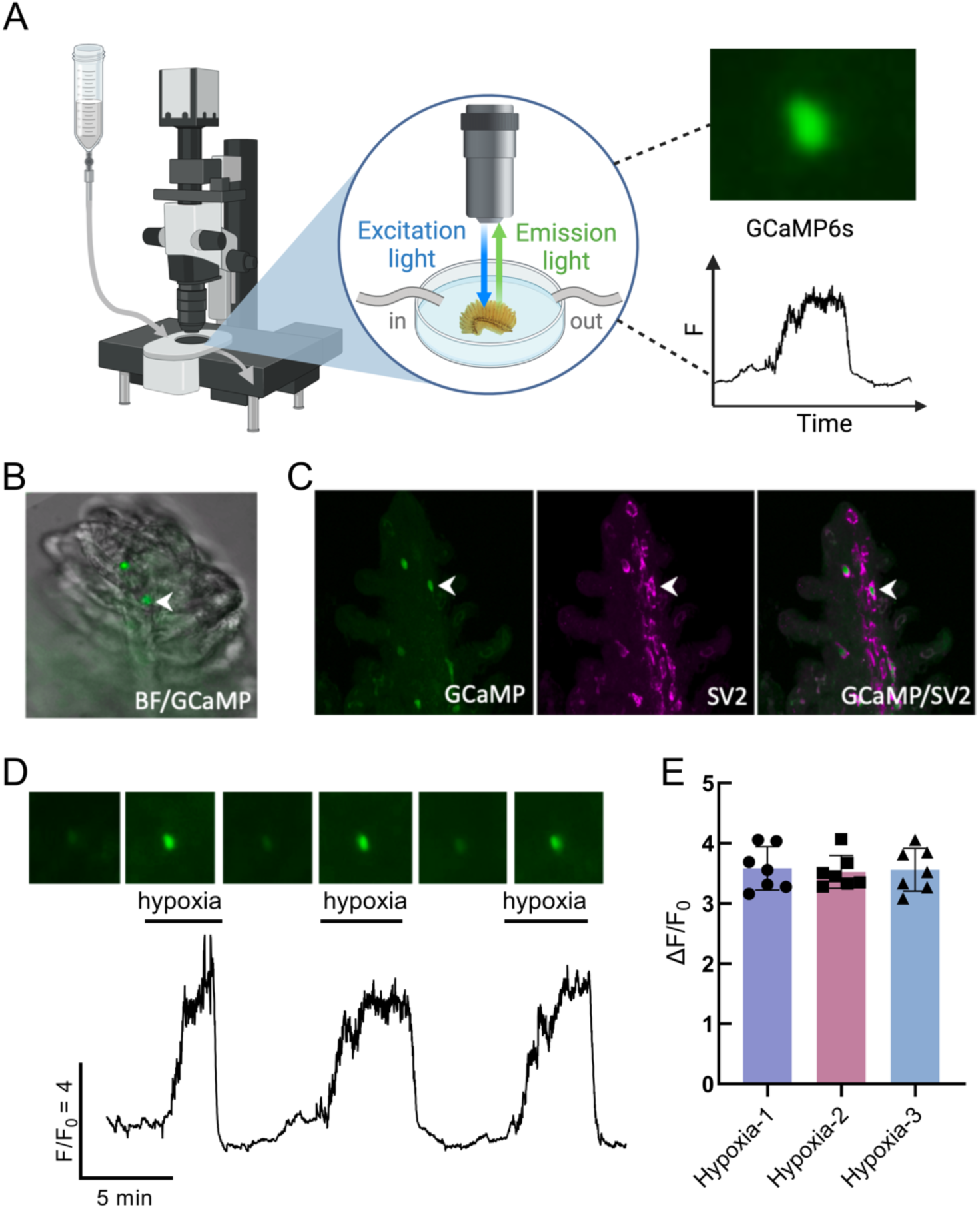
Hypoxia induced intracellular Ca^2+^ responses in gill neuroepithelial cells from Tg(*elavl3*:GCaMP6s) zebrafish. (A) Schematic of the GCaMP recording preparation illustrating (left) a fluorescence microscope, (centre) an isolated gill in a recording chamber fitted with in-and out-flow for superfusion, and (right) fluorescence excitation in a cell during a hypoxic stimulus. (B) Overlay of brightfield and green fluorescence (488 nm) images of a GCaMP-positive neuroepithelial cell (NEC, arrowhead) *in situ* containing GCaMP from a Tg(*elavl3:*GCaMP6s) zebrafish. (C) Post hoc confocal imaging confirming immunohistochemical co-localization of GCaMP (green) with synaptic vesicle protein-2 (SV2, magenta) in the NEC (arrowheads) identified in (B). The panels show (from left) GCaMP and SV2 separately, and GCaMP and SV2 labelling together. (D) Ca^2+^ imaging trace from the GCaMP-containing cell in (A) during three bouts of hypoxia. Scale indicates time (min) and relative changes in fluorescence (F/F_0_) corresponding to changes in intracellular Ca^2+^ concentration ([Ca^2+^]_i_). Time-series micrographs above show fluorescent changes over time. (E) Average (±S.E.M.) F/F_0_ in NECs in response to three consecutive bouts of hypoxia. There was no significant change in the magnitude of the Ca^2+^ response to hypoxia over time (Kruskal-Wallis test, P>0.999, n=7). Panel A created with BioRender.com.

### Extracellular and stored Ca^2+^ contribute to the NEC response to hypoxia

A combination of intracellular and extracellular blockers were used to evaluate the source of Ca^2+^ in the NEC response to hypoxia. In experiments where NECs were exposed to successive bouts of hypoxia, the Ca^2+^ response was significantly reduced by 50.4% with the addition of 100 μM nifedipine, an L-type Ca^2+^ channel blocker (Fig. 3A). The Ca^2+^ response fully recovered after 15 min washout of nifedipine (Fig. 3A,B; Kruskal-Wallis test, P=0.006, n=6). Similarly, the response to hypoxia was partially reduced by 41.9% when Ca^2+^ was removed from the extracellular solution (Fig. 2C,D; Kruskal-Wallis test, P<0.0059, n=6).

**Figure 3.**
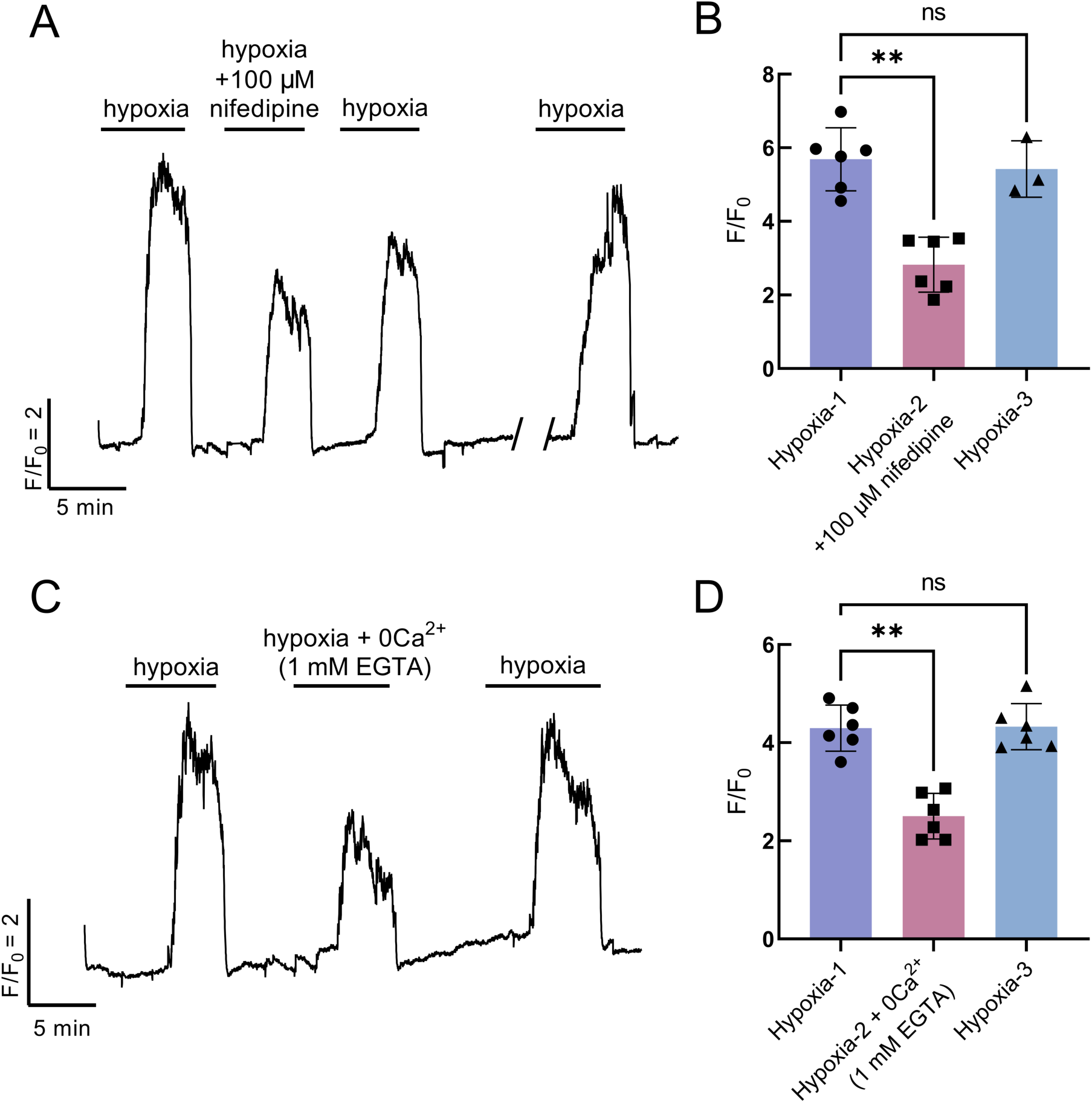
Extracellular Ca^2+^ contributes to the response to hypoxia in NECs. (A) Ca^2+^ imaging trace from a GCaMP-containing NEC, where the response to hypoxia was reduced with the addition of 100 µM nifedipine, an L-type Ca^2+^ channel blocker. The response to hypoxia was fully recovered after 15 min washout (break in trace). (B) Summary data from NECs as treated in (A) illustrating a reduction in the average (±S.E.M.) F/F_0_ (Kruskal-Wallis test, P=0.006, n=6). 3 cells were evaluated for recovery after a 15-min washout period (Kruskal-Wallis test, P>0.99, n=3). (C) Ca^2+^ imaging trace from a GCaMP-containing NEC, where the response to hypoxia was reversibly reduced when Ca^2+^ was removed from the extracellular solution. (D) Summary data from NECs as treated in (C) showing a reduction in the average (±S.E.M.) F/F_0_ (Kruskal-Wallis test, P<0.0059, n=6). The response to hypoxia fully recovered after zero Ca^2+^ treatment (Kruskal-Wallis test, P>0.99).

The NEC Ca^2+^ response to hypoxia was partially reduced by 30.4% with the addition of dantrolene, an inhibitor of intracellular Ca^2+^ release (Fig. 4A,C; Kruskal-Wallis test, P=0.007, n=6). To confirm that the Ca^2+^ response to hypoxia was a reflection of the sum of Ca^2+^ arising from both intracellular and extracellular sources, we exposed NECs to hypoxia in the presence of dantrolene and Ca^2+^-free solution (Fig. 4B,D; Kruskal-Wallis test, P<0.0079, n=5). When compared to all other treatments, blocking both intracellular and extracellular Ca^2+^ resulted in the largest reduction in the Ca^2+^ response to hypoxia (Fig. 4E).

**Figure 4.**
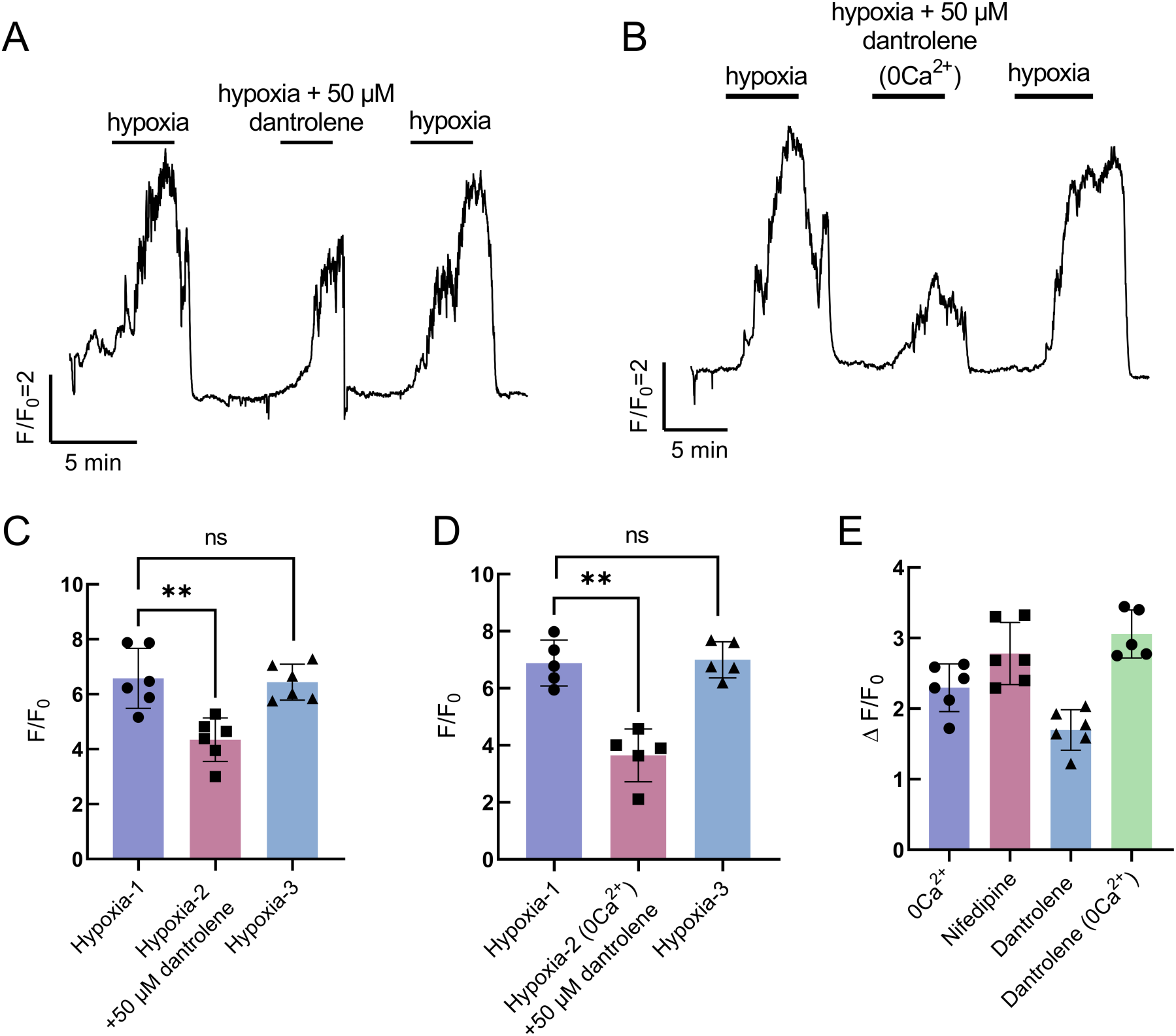
Intracellular Ca^2+^ contributes to the response to hypoxia in NECs. (A) Ca^2+^ imaging trace from a GCaMP-containing NEC, where the response to hypoxia was reversibly reduced with the addition of 50 µM dantrolene, an inhibitor of intracellular Ca^2+^ release. (B) Ca^2+^ imaging trace from a GCaMP-containing NEC demonstrating the combined contributions of intracellular and extracellular Ca^2+^. The response to hypoxia was further reduced with the addition of 50 µM dantrolene in Ca^2+^-free extracellular solution. (C) Summary data as treated in (A) showing reduction in average (±S.E.M.) F/F_0_ (Kruskal-Wallis test, P=0.007, n=6). The response to hypoxia fully recovered (Kruskal-Wallis test, P>0.99, n=6). (D) Summary data as treated in (B) showing a reduction in average (±S.E.M.) F/F_0_ (Kruskal-Wallis test, P<0.0079, n=5). The response to hypoxia fully recovered (Kruskal-Wallis test, P>0.99, n=5). (E) Summary comparing all Ca^2+^ blocking treatments. Blocking both intracellular and extracellular Ca^2+^ resulted in the largest change in the Ca^2+^ response to hypoxia (ΔF/F_0_).

### The NEC Ca^2+^ response to hypoxia is reduced by D_2_ receptor activity

Previous studies reported a decrease in ventilation frequency of larval zebrafish with dopamine or the specific dopamine D_2_ receptor agonist, quinpirole (Shakarchi et al., 2013; Reed et al., 2024). Additionally, immunohistochemical labelling has shown co-localization of dopamine D_2_ receptors with NECs and has provided evidence for the postsynaptic synthesis and reuptake of dopamine in the gill (Reed et al., 2024). In this study, we sought to determine an inhibitory role for D_2_ receptors within the gill at the level of the presynaptic NEC. We found the NEC Ca^2+^ response to hypoxia was reversibly reduced by 44.1% with the addition of dopamine (Fig. 5A,B; Kruskal-Wallis test, P=0.0116, n=5), as well as reduced by 57.9% with the addition of quinpirole (Figure 5C,D; Kruskal-Wallis test, P=0.007, n=6). Domperidone, a D_2_ receptor antagonist, had the opposite effect and enhanced the NEC Ca^2+^ response to hypoxia by 26.7% compared to hypoxia alone (Fig. 5E,F; Kruskal-Wallis test, P=0.0347, n=6).

**Figure 5.**
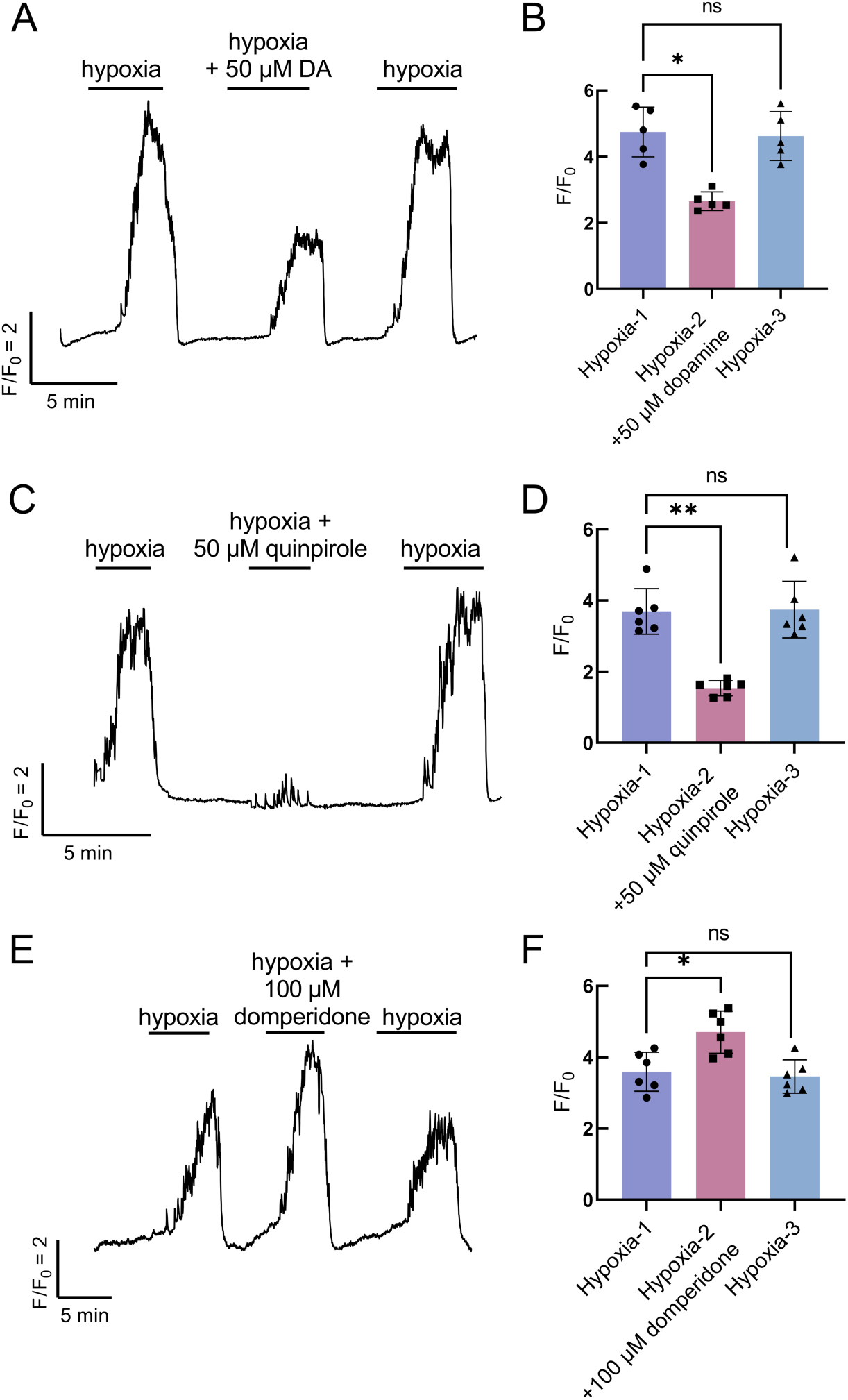
The effects of dopamine and D_2_ receptor activity on the NEC response to hypoxia. (A) Ca^2+^ imaging trace from a GCaMP-containing NEC where the response to hypoxia was reversibly reduced with the addition of 50 µM dopamine (DA). (C) Ca^2+^ imaging trace from a GCaMP-containing NEC where the response to hypoxia was reversibly reduced with the addition of 50 µM quinpirole, a specific dopamine D_2_-receptor agonist. (E) Ca^2+^ imaging trace from a GCaMP-containing NEC where the response to hypoxia was enhanced with the addition of 100 µM domperidone, a specific dopamine D_2_-receptor antagonist. (B,D,F) Summary data showing average (±S.E.M.) F/F_0_ corresponding to experiments in (A,C,E). Addition of 50 µM dopamine significantly reduced the Ca^2+^ response to hypoxia (B, Kruskal-Wallis test, P=0.0116, n=5), as well as 50 µM quinpirole (D, Kruskal-Wallis test, P=0.007, n=6), while domperidone increased average (±S.E.M.) F/F_0_ (F, Kruskal-Wallis test, P=0.0347, n=6). The response to hypoxia fully recovered following all treatments (Kruskal-Wallis test, P>0.99, n=5-6).

To evaluate an intracellular pathway for the inhibition of the NEC Ca^2+^ response to hypoxia via dopamine D_2_ receptors, we used drugs to target AC. Activation of D_2_ reduces AC activity (Usiello et al., 2000). We found a decrease in the response to hypoxia with the addition of SQ22536, an inhibitor of AC (Fig. 6A,B; Kruskal-Wallis test, P=0.0099, n=6). Further, when the Ca^2+^ response to hypoxia was reduced in the presence of dopamine, activation of AC with forskolin partially restored the normal Ca^2+^ response to hypoxia (Fig. 6C,D; Kruskal-Wallis test, P=0.0019, n=6).

**Figure 6.**
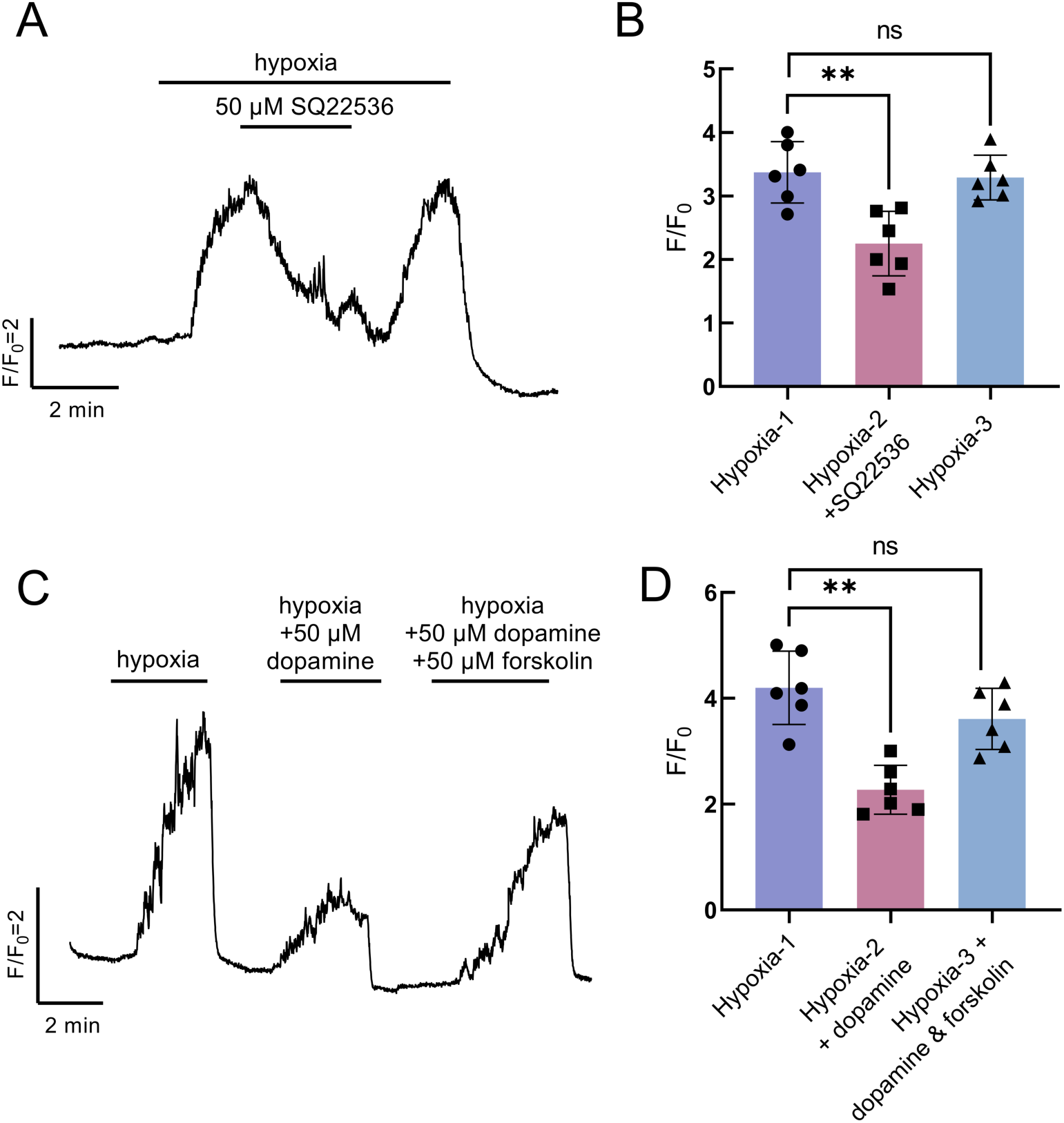
Dopamine acts through intracellular secondary messenger cyclic adenosine 3′,5′-monophosphate (cAMP) in NECs. (A) Addition of SQ22536, an adenylyl cyclase (AC) inhibitor, decreased the effect of hypoxia on intracellular Ca^2+^. (B) Summary data as treated in (A) showing reduction in average (±S.E.M.) F/F_0_ (Kruskal-Wallis test, P=0.0099, n=6). (C) Forskolin, an AC activator, partially recovered the suppressive effect of dopamine on the Ca^2+^ response to hypoxia. (D) Summary data as treated in (C) showing recovery in average (±S.E.M.) F/F_0_ of the hypoxic response from dopamine with forskolin (Kruskal-Wallis test, P=0.0019, n=6). The response to hypoxia fully recovered following both treatments (Kruskal-Wallis test, P>0.99, n=6).

### Postsynaptic responses to hypoxia are modulated by presynaptic D_2_ activity

In addition to NECs, immunohistochemical labelling confirmed that SV2/zn-12-positive ChNs in the gill of Tg(*elavl3*:GCaMP6s) zebrafish are GCaMP-positive (Fig. 7). ChNs extend the complete length of gill filaments and were previously shown to innervate NECs in zebrafish (Jonz and Nurse, 2003). Rotation of confocal images by 90° shows zn-12-positive nerve fibres in close association with GCaMP-positive NECs and ChNs (Fig. 7D-I).

**Figure 7.**
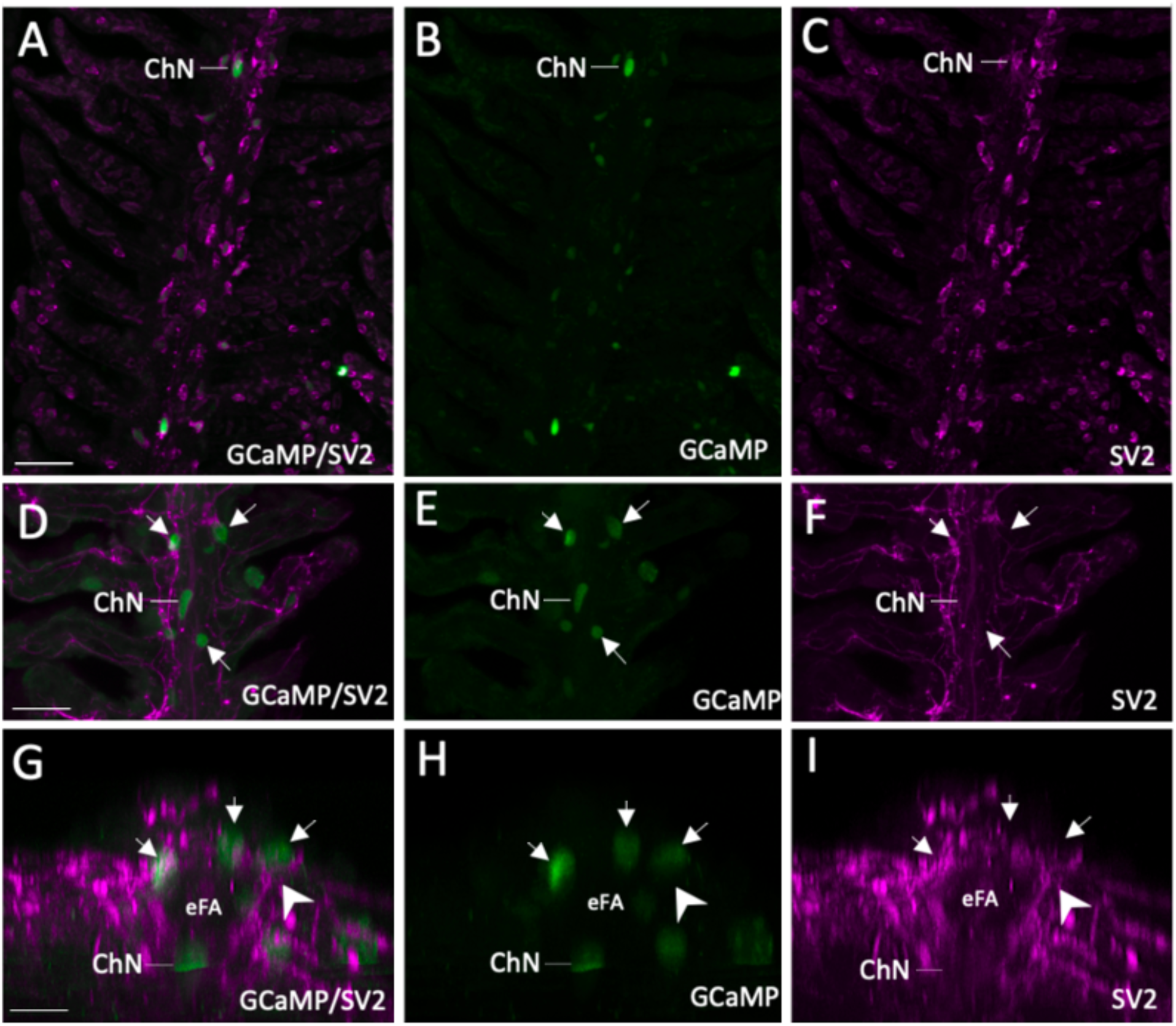
Characterization of GCaMP-positive postsynaptic chain neurons in Tg(*elavl3*:GCaMP6s) zebrafish gills. (A-C) confocal imaging of immunohistochemical localization of GCaMP in a chain neuron (ChN) containing synaptic vesicle protein-2 (SV2, magenta). (B,C) GCaMP and SV2 labelling shown separately. Scale bar in A=50 µm and applies to B and C. (D-F) co-labelling of GCaMP-positive ChNs and NECs (green, arrows) with nerve fibres labelled with zn-12 (magenta). (E, F) GCaMP and zn-12 labelling shown separately. Scale bar in D=50 µm and applies to E and F. (G-I) Images from panels (D-F) cropped and titled back 90°. Rotation demonstrates neural connection (arrowhead) between nerve fibres located below the efferent filament artery (eFA) where ChNs are located, projecting to GCaMP-positive NECs.

In our whole-gill recording preparation, multiple ChNs of a single gill filament showed nearly simultaneous Ca^2+^ responses to hypoxia (Fig. 8A,B). Since NECs are confined to the distal end of the gill filament, the filament was transected at a proximal region where ChNs were present but no NECs were observable (Fig. 8C). This technique was used to mechanically remove synaptic contact between NECs and ChNs to determine whether ChNs can respond to hypoxia independently of NECs. After the filament was cut, the ChN response to hypoxia was completely abolished, though ChNs were still able to show a response to high K^+^, indicating that ChNs were not adversely affected by the cut (Fig. 8D). To further confirm NECs were innervated by ChNs, fish were treated with 6-OHDA, a neurotoxin used to destroy the nerve terminals of dopaminergic neurons. Since NECs are innervated by nerve fibres containing the dopamine active transporter (DAT) (Reed et al., 2023), this technique was used to chemically remove the ability of postsynaptic ChNs to receive inputs from NECs. In transgenic animals produced by crossing the Tg(*dat:tom20 MLS-mCherry*) and Tg(*elavl3*:GCaMP6s) lines, labelling of DAT-positive ChN terminals around NECs was partially reduced in fish treated with 6-OHDA (Fig. 8E). Importantly, after treatment with 6-OHDA, ChNs showed no response to hypoxia while still maintaining the ability to respond to high K^+^ (Fig. 8F, overlayed traces, n=6).

**Figure 8.**
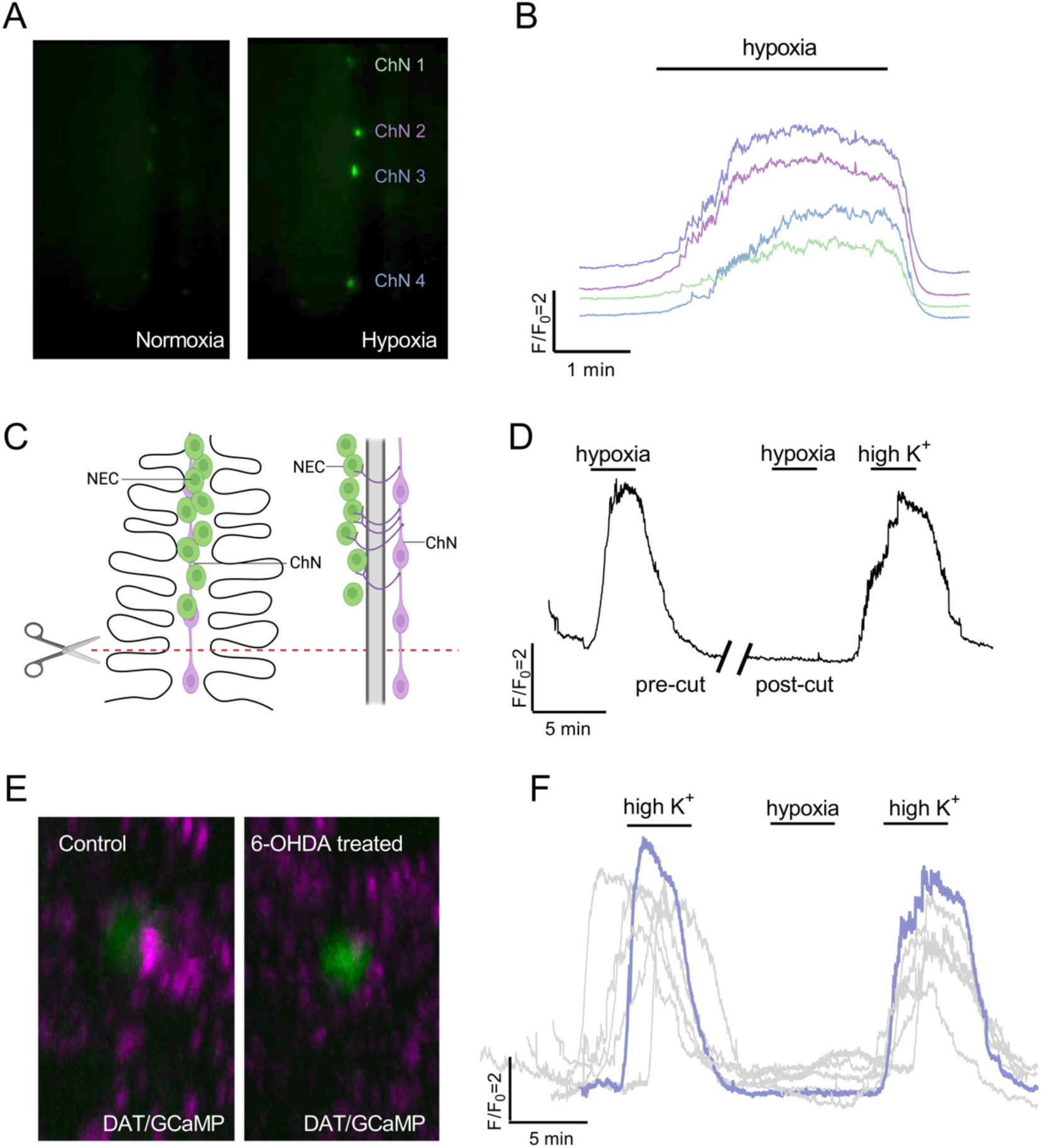
The chain neuron (ChN) calcium response to hypoxia requires synaptic contact with NECs. (A) Examples from live fluorescence imaging at 488 nm of four ChNs along a single filament in normoxia (left micrograph) and hypoxia (right micrograph). (B) Calcium traces from the four ChNs shown in (A) responding to hypoxia. Different colours represent corresponding cells (ChN 1-4) in (A). (C) Schematic of a gill filament (left) and rotation by 90° on the y-axis (right) illustrate filament transection. Filaments were cut along the red dashed line at the proximal end where a ChN was present, but no NECs were observable. (D) Calcium imaging trace of a single ChN before and after filament transection. After the filament was cut (break in trace), the neuron no longer responded to hypoxia. As a positive control, viability of the neuron was demonstrated by stimulation with a solution of high extracellular K^+^. (E) Confocal imaging of gills from a double transgenic animal produced by crossing Tg(*elavl3*:GCaMP6s) and Tg(*dat:tom20 MLS-mCherry*) fish showing the relationship between the dopamine active transporter (DAT) nerve endings (magenta) and GCaMP-positive NECs (green) in control (left micrograph) and 6-OHDA-treated gills (right micrograph). DAT labelling was found in close proximity to NECs but was reduced after 6-OHDA treatment. (F) Overlayed Ca^2+^ imaging traces from 6-OHDA treated animals (n=6). After 6-OHDA treatment the ChNs did not respond to hypoxia. Response to high extracellular K^+^ confirmed cell viability. Panel C created with BioRender.com.

Finally, we aimed to link our characterization of D_2_ receptor activity in NECs with the Ca^2+^ response in postsynaptic ChNs. High K^+^ was used to depolarize the membranes of ChNs with and without the addition of quinpirole. In agreement with the reported absence of D_2_ receptors on ChNs (Reed et al., 2023), both 50 µM and 100 µM quinpirole failed to reduce the [Ca^2+^]_i_ response during a high K^+^ stimulus, confirming that dopamine and quinpirole do not affect ChNs directly and must be acting presynaptically (Fig. 9A,B). In dual recording of NECs and ChNs, in which synaptic contact between cells was left intact, [Ca^2+^]_i_ was simultaneously recorded in both cell types during exposed to hypoxia (Fig. 9C). Under these conditions, NECs responded first with an increase in [Ca^2+^]_i_, followed by a [Ca^2+^]_i_ response in ChNs after a latency of 20-30 s (Fig. 9D). There was a reduction in both the NEC and ChN Ca^2+^ response to hypoxia with the D_2_ receptor agonist, quinpirole (Fig. 9D,E; NEC: Kruskal-Wallis test, P=0.003, n=5, and ChN: Kruskal-Wallis test, P=0.013, n=5).

**Figure 9.**
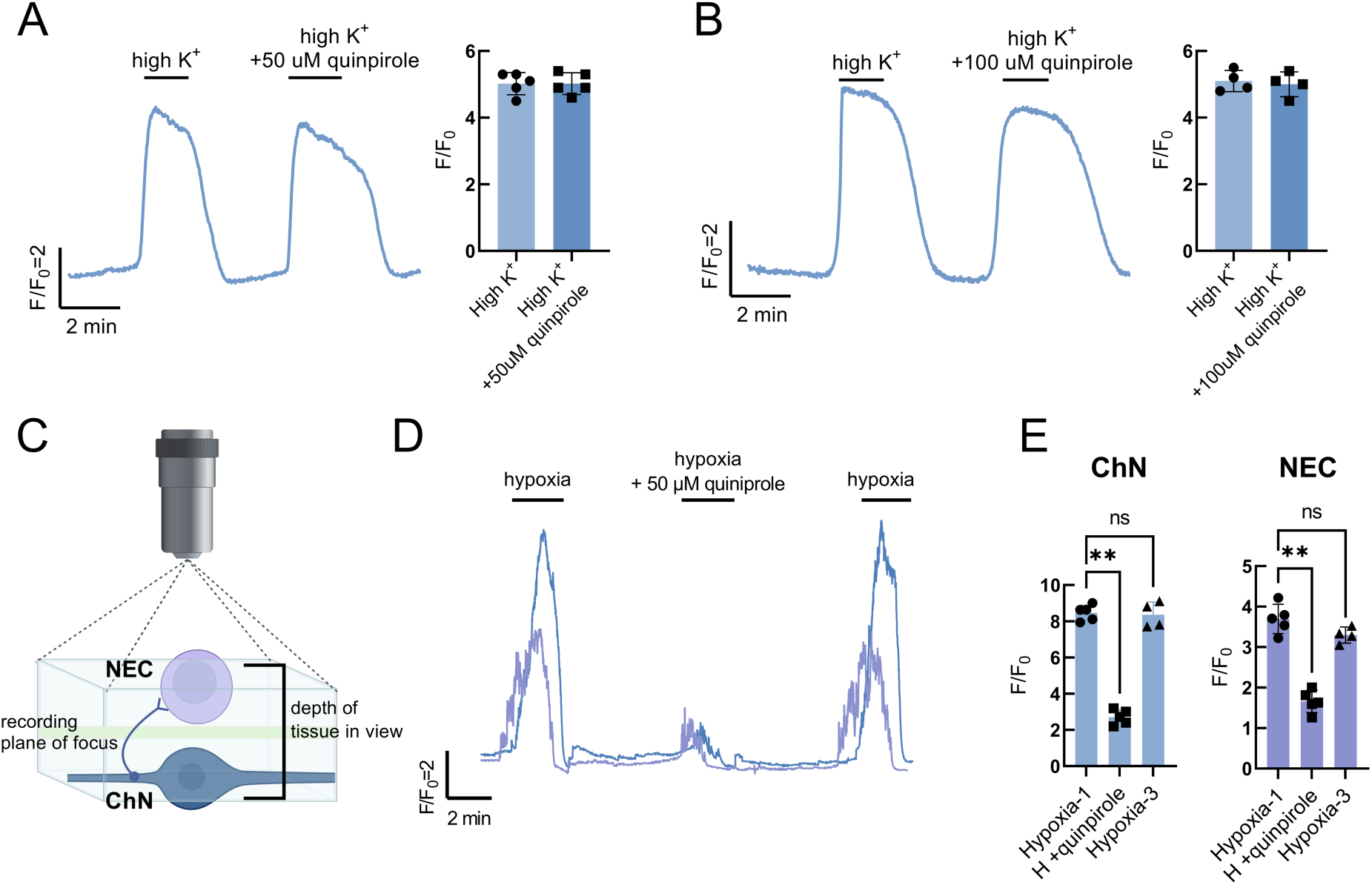
Postsynaptic modulation of the hypoxic response by presynaptic D_2_ receptor activation. (A) Calcium imaging trace from a single chain neuron with summary data showing no change in the response to high K^+^ with the addition of 50 µM quinpirole. (B) Calcium imaging trace from a single chain neuron (ChN) with summary data showing no change in the response to high K^+^ with the addition of 100 µM quinpirole. (C) Schematic illustration of preparation for dual recording of NEC and ChN. Focus plane for recording was set at a tissue depth between both cells (green line) where both cells were still in view (blue box). (D) Dual-recording Ca^2+^ imaging trace of a NEC (purple) and ChN (blue) recorded simultaneously. (E) Summary data as treated in D showing a decrease in the hypoxic signal produced by NECs (Kruskal-Wallis test, p=0.003, n=5) and ChNs (Kruskal-Wallis test, p=0.013, n=5) with quinpirole. In both NECs and ChNs, the hypoxic response was fully recovered after quinpirole treatment (Kruskal-Wallis test, p>0.999, n=4). Panel C created with BioRender.com

## Discussion

The present study demonstrates neuromodulation of the chemoreceptor response to hypoxia produced by activation of D_2_ receptors in NECs in the gills of zebrafish. We used a transgenic zebrafish line in which the Ca^2+^ reporter, GCaMP, was genetically encoded to reveal dopaminergic inhibition of the NEC Ca^2+^ response to hypoxia, acting through D_2_ and inhibition of AC. We further demonstrated a link between the inhibitory effect of activation of presynaptic D_2_ and modulation of the hypoxic signal in postsynaptic neurons that innervate NECs.

### The gill as a model for the chemoreceptor response to hypoxia

Evaluating responses of chemoreceptors to hypoxia in previous work has typically relied on the techniques of classical Ca^2+^ imaging, where the loading of Ca^2+^ reporting dyes into isolated cells was required in the carotid body (Montoro et al. 1996), neuroepithelial bodies (Livermore et al., 2015) and gill NECs (Zhang et al., 2011; Porteus et al., 2014; Abdallah et al., 2015; Leonard et al., 2022). Here, we developed an intact, whole-gill preparation using the endogenous Ca^2+^ indicator, GCaMP. This preparation allows us to observe chemoreceptor responses to hypoxia without the added complication of dye loading, while maintaining functional connectivity throughout the organ. Gill GCaMP-positive chemoreceptors showed a reliable response to hypoxia that was consistent over time and multiple hypoxic stimuli, serving as a useful model to evaluate chemoreceptor responses to hypoxia.

Though several studies have used Ca^2+^ imaging to evaluate chemoreceptor activity in gill NECs (Zhang et al., 2011; Porteus et al., 2014; Abdallah et al., 2015; Leonard et al., 2022), the source of Ca^2+^ contributing to the response to hypoxia in NECs had not clearly been determined. In the present study, we determine the NEC Ca^2+^ response to hypoxia involves both extracellular Ca^2+^ influx through L-type Ca^2+^ channels, as well as release from intracellular stores.

### Dopaminergic modulation of chemoreceptor activity during hypoxia

In the present study, we demonstrated a reduction in the NEC Ca^2+^ response to hypoxia produced by application of dopamine or the specific D_2_ receptor agonist, quinpirole. D_2_ receptors were previously localized to gill NECs in zebrafish by immunohistochemistry, whereas D_2_ was not found on postsynaptic ChNs (Reed et al., 2024). In addition, D_2_ expression in NECs was demonstrated using single-cell RNA-sequencing (Pan et al., 2022). Accordingly, the present results confirm that ChNs showed no direct response to quinpirole. We therefore attribute the suppressive effects of dopamine and quinpirole to activation of presynaptic D_2_ receptors.

Dopamine receptors are a type of G-protein-coupled receptor (GPCR) that play a crucial role in mediating the effects of the neurotransmitter, dopamine. There are five known subtypes of dopamine receptors classified into two main families based on their signaling mechanisms: D_1_-like (D_1_ and D_5_), and D_2_-like (D_2_, D_3_, and D_4_). Upon dopamine binding, dopamine receptors undergo conformational changes that activate intracellular signaling pathways through interactions with G-proteins. Activation of D_2_-like receptors inhibits AC activity through Gαi/o proteins, leading to decreased cAMP levels within the cell (reviewed by Beaulieu & Gainetdinov, 2011). We showed that inhibition of AC with SQ22536 mimicked the modulation of the NEC hypoxic response by dopamine. Further, the modulatory effects of dopamine were reversed by addition of forskolin, an AC activator. Together, these results provide evidence for the dopaminergic inhibition of AC, and suggest subsequent decreases in cAMP activity, leading to modulation of the NEC Ca^2+^ response to hypoxia. This mechanism of inhibition by D_2_ receptors is similar to carotid body type 1 cells, where dopamine causes a dose-dependent decrease in the type 1 cell cAMP content induced by forskolin (Batuca et al., 2003)

### Dopaminergic modulation of gill hypoxia signalling

The present study demonstrates modulation of the ChN response to hypoxia via presynaptic D_2_ receptor activation, which suggests D_2_ at the NEC-ChN synapse may act on ventilation. ChNs are part of a nerve bundle in the gill located beneath the efferent filament artery that sends projections to innervate NECs. Using gill transection experiments we identified a pathway for the transmission of the hypoxic signal in the gill through a functional unit containing two parts: the postsynaptic ChN, and any number NECs innervated by that ChN. We further showed that within this functional unit, the inhibitory actions of dopamine on the presynaptic NEC can be observed in the postsynaptic ChN, suggesting a mechanism of modulation at the NEC-ChN synapse. This mechanism may be similar to the carotid body, where dopamine released by type 1 cells has an autocrine-paracrine action on dopaminergic D_2_-receptors located on type 1 cells to inhibit Ca^2+^-channels, leading to negative feedback regulation of further neurotransmitter release during hypoxia (Benot & López-Barneo, 1990). However, the excitatory neurotransmitter in the NEC-ChN synapse has yet to be elucidated.

ChNs were previously shown to innervation NECs in gills of zebrafish (Jonz & Nurse, 2003), but their physiological role has been elusive. We propose that ChNs may act as interneurons that carry a hypoxic signal from NECs to afferent fibres of the glossopharyngeal or vagus nerves−the cranial nerves that carry chemoreceptor activity to the CNS in fish (Burleson & Milsom, 1995; Sundin & Nilsson, 2002). ChNs contain varicose processes that indicate multiple locations of synaptic contact, possibly with the adjacent sensory fibres of the extrinsic nerve bundle (Jonz and Nurse, 2003). Furthermore, in zebrafish larvae exogenous application of dopamine reduces ventilation frequency, whereas the D_2_ agonist, quinpirole, attenuates the hyperventilatory response to hypoxia (Shakarhi et al., 2013; Reed et al., 2024). These results, combined with the present findings, suggest the observed decrease in hyperventilation is the result of presynaptic D_2_ activation and a reduction in NEC activity.

Intriguingly, in the present study we report that the D_2_ receptor antagonist, domperidone, enhanced the NEC Ca^2+^ response to hypoxia. This observation points to endogenous dopamine release during hypoxia acting upon presynaptic D_2_ receptors to limit the cellular response to hypoxia. Previous work has identified postsynaptic neurons, including ChNs, as a source of dopamine production and storage in the gill (Reed et al., 2024). Presynaptic type 1 cells are largely responsible for synthesizing and releasing dopamine in the mammalian carotid body. However, many carotid body afferents are also dopaminergic (Finley et al., 1992), and carotid sinus nerve fibres innervating type 1 cells release dopamine (Almaraz et al., 1993), suggesting an additional postsynaptic source of dopamine to further modulate the type 1 cell response to hypoxia, similar to that observed in the gill.

### Implications

The results of this study have implications for the field of oxygen sensing, and may contribute to our understanding of acclimatization, development, and evolution in vertebrates. For example, modulation of presynaptic chemoreceptor inhibition by D_2_ may contribute to hypoxia acclimatization. In zebrafish, total gill D_2_ receptor mRNA expression decreases after 48 h of hypoxia exposure, a timepoint in acclimation where there is a shift away from aquatic surface respiration behaviour, allowing the animal to rely more heavily on hyperventilation (Reed et al., 2024). Acclimation may lower the number of inhibitory D_2_ receptors expressed by NECs and lead to a change in chemoreceptor sensitivity of the cell to dopamine. A similar involvement of D_2_ receptors in hypoxia acclimation has been proposed in the carotid body by Huey and Powell (2000), who showed decreased carotid body D_2_ receptor mRNA expression following hypoxia acclimation and suggested this may be one mechanism by which exposure to chronic hypoxia reduces inhibition of hypoxia signaling to enhance ventilatory acclimatization to hypoxia.

The zebrafish model used in this study may be particularly useful in characterizing the mechanisms involved in age-related changes in chemosensitivity. In the mammalian carotid body, expression of genes encoding important components of the dopaminergic system are altered during development. These changes coincide with changes in chemoreceptor activity and output observed in early postnatal development in rats (Gauda et al., 1996, Gauda 2002, Bairam & Carroll 2005). Zebrafish have already shown an adaptable dopaminergic system under hypoxia (Reed et al., 2024), and thus may serve as a great model for exploring changes in chemoreceptor sensitivity during development.

D_2_ is the predominant dopamine receptor in the carotid body, which is consistent with what we have found here in the zebrafish gill; however, that does not rule out contributions of other dopamine receptors such as D_1_. mRNA expression of D_1_ receptors has been found in the carotid body of rats, cats, and rabbits (Bairam et al., 1998). Further, RNA sequencing showed expression of other dopamine receptors in the gill, specifically D_1_-like receptors in NECs (Pan et al., 2022). Zebrafish have served as an excellent model to explore the evolutionary origins of dopamine, and D_2_ receptors, and may be particularly useful in clarifying a role for other dopamine receptors in hypoxia signalling.

## Conclusion

The present study delineated a mechanism by which presynaptic dopamine D_2_ receptors provide a feedback mechanism that attenuates the chemoreceptor response to hypoxia. We found activation of presynaptic D_2_ receptors decreases the NEC Ca^2+^ response to hypoxia via intracellular AC inhibition. We provide evidence for postsynaptic modulation of the Ca^2+^ response to hypoxia via presynaptic D_2_ receptors and suggest a link between chemoreceptor inhibition by dopamine and modulation of the hypoxic ventilatory response. Our results provide the first evidence of neuromodulation of the hypoxic signal produced by NECs in the gill and suggests that a modulatory role for dopamine in oxygen sensing arose early in vertebrate evolution.

## Additional information

### Data availability statement

All data supporting the results are presented within the figures. Raw data may additionally be made available upon request.

### Author contributions

M.R. and M.G.J. conceived of and designed the experimental approach, and were involved in interpretation of the results. M.R. performed the experiments, data analysis, preparation of the figures, and wrote the manuscript. M.R. and M.G.J. edited and revised the manuscript. Both authors approve the final version of this manuscript and are accountable for all aspects of the work.

